# QSIPrep: An integrative platform for preprocessing and reconstructing diffusion MRI

**DOI:** 10.1101/2020.09.04.282269

**Authors:** Matthew Cieslak, Philip A. Cook, Xiaosong He, Fang-Cheng Yeh, Thijs Dhollander, Azeez Adebimpe, Geoffrey K. Aguirre, Danielle S. Bassett, Richard F. Betzel, Josiane Bourque, Laura M. Cabral, Christos Davatzikos, John Detre, Eric Earl, Mark A. Elliott, Shreyas Fadnavis, Damien A. Fair, Will Foran, Panagiotis Fotiadis, Eleftherios Garyfallidis, Barry Giesbrecht, Ruben C. Gur, Raquel E. Gur, Max Kelz, Anisha Keshavan, Bart S. Larsen, Beatriz Luna, Allyson P. Mackey, Michael Milham, Desmond J. Oathes, Anders Perrone, Adam R. Pines, David R. Roalf, Adam Richie-Halford, Ariel Rokem, Valerie J. Sydnor, Tinashe M. Tapera, Ursula A. Tooley, Jean M. Vettel, Jason D. Yeatman, Scott T. Grafton, Theodore D. Satterthwaite

## Abstract

Diffusion-weighted magnetic resonance imaging (dMRI) has become the primary method for non-invasively studying the organization of white matter in the human brain. While many dMRI acquisition sequences have been developed, they all sample q-space in order to characterize water diffusion. Numerous software platforms have been developed for processing dMRI data, but most work on only a subset of sampling schemes or implement only parts of the processing workflow. Reproducible research and comparisons across dMRI methods are hindered by incompatible software, diverse file formats, and inconsistent naming conventions. Here we introduce QSIPrep, an integrative software platform for the processing of diffusion images that is compatible with nearly all dMRI sampling schemes. Drawing upon a diverse set of software suites to capitalize upon their complementary strengths, QSIPrep automatically applies best practices for dMRI preprocessing, including denoising, distortion correction, head motion correction, coregistration, and spatial normalization. Throughout, QSIPrep provides both visual and quantitative measures of data quality as well as “glass-box” methods reporting. Taken together, these features facilitate easy implementation of best practices for processing of diffusion images while simultaneously ensuring reproducibility.

## INTRODUCTION

The computations in the human brain that give rise to cognition and behavior rely in part on spatially distributed regions that are connected via myelinated axons. Water diffusion is hindered and restricted in the presence of these myelinated axons, allowing axonal organization to be characterized using diffusion-weighted MRI (dMRI). As the technology underlying dMRI has rapidly advanced, it has become the primary technique for non-invasive studies of white matter organization in humans. dMRI typically samples three spatial dimensions and three additional dimensions called “*q*-space,” measuring signals that reflect the water diffusion process at a spatial location, along a given direction, with a specified sensitivity. The diffusion process can be mathematically characterized (“reconstructed”) in each voxel, such that physical properties of local white matter microstructure can be estimated based on variability in the diffusion process. As a method, dMRI has benefitted from dramatic improvements in *q*-space sampling strategies^1,2^, methods to estimate diffusion propagators^2–4^ and approaches to more accurately relate observed MR signal to white matter structures^5–8^ (see **Supplementary Note 1** for an overview).

However, this rapid progress has come with costs. Both the increased diversity of *q-* space sampling schemes used in image acquisition and the proliferation of analysis packages to process and reconstruct dMRI data are a cause of confusion among translational investigators. Two major obstacles are evident. First, many pre-processing and reconstruction methods are dependent on specific *q*-space sampling schemes. Second, even within a given sampling scheme, it is difficult for researchers to move between methods implemented by different software packages, due to the disparate naming conventions and file formats used. Even when an investigator does manage to implement more than one analysis workflow, it is often challenging to compare them due to the divergent measures produced by different software analysis packages. As a result, most labs tend to use a limited set of software packages, failing to capitalize upon the complementary capabilities of different tools.

In response to these obstacles, we introduce QSIPrep, a unified and robust platform for processing and reconstructing dMRI data. QSIPrep leverages the metadata recorded in the Brain Imaging Data Structure (BIDS)^9^, a widely used specification that provides a standardized means for describing imaging data. QSIPrep automatically builds preprocessing pipelines by using BIDS to detect critical scanning parameters such as the *q*-space sampling scheme, phase encoding (PE) direction, total readout time, and fieldmap configurations. Processing pipelines are then automatically configured based on the available data, allowing adaptive preprocessing pipelines to accommodate diverse *q*-space sampling schemes. The pipelines generated by QSIPrep are fully documented by visual reports at each step as well as standardized text that provides a methodological description of the pipeline. All software dependencies are included in a freely available Docker image that is subject to continuous integration testing, allowing proven software to be executed in nearly any environment.

In addition to preprocessing, QSIPrep implements advanced reconstruction and tractography methods in curated reconstruction workflows using tools from leading software packages. Post-processing methods from DSI Studio^10^, DIPY^11^ and MRtrix^12^ are encapsulated in self-documented workflows that consume the output from QSIPrep’s preprocessing pipeline. Critically, as each of these software packages uses different conventions and file formats, QSIPrep reconstruction workflows include a final step in which all output – including reconstructions, scalar maps, tractograms, and connectivity matrices – are converted to a consistent, interoperable format. Together, QSIPrep provides a framework for uniform workflows that support both the diversity of dMRI sampling schemes and state-of-the-art analytic approaches. As a result, QSIPrep allows translational scientists to easily implement best practices for dMRI in a fully reproducible manner.

## RESULTS

QSIPrep has been publicly available since December 2019. At the time of writing, it has been downloaded over 2,200 times and has been used to process over 21,000 dMRI scans according to our application and error tracking system. Continuous integration testing and the open development environment enable rapid bug detection and feature requests from an international user base. Below, we demonstrate that QSIPrep automatically generates preprocessing workflows for all static *q*-space sampling methods. These sampling methods include spherical sampling (both single-shell and multi-shell), Cartesian grid sampling (sometimes referred to as Diffusion Spectrum Imaging; DSI), and random sampling. Notably, we demonstrate that QSIPrep performs comparably or better than published workflows that were explicitly tailored for each sampling scheme. Finally, we illustrate how preprocessed images can be reconstructed using QSIPrep’s diverse set of curated reconstruction workflows, yielding results that are conveniently visualized and stored in a common file format.

### A BIDS app that builds appropriate preprocessing workflows

QSIPrep began as a fork of the popular fMRIPrep software^13^, building on its use of modular and adaptive workflows. As for fMRI, there are numerous software packages available to process and reconstruct dMRI data. QSIPrep uses software from FSL^14^, MRtrix3^12^, DIPY^11^, DSI Studio^10^ and ANTs^15^, among others. **Figure 1** illustrates QSIPrep’s preprocessing and reconstruction workflows.

**Fig 1.**
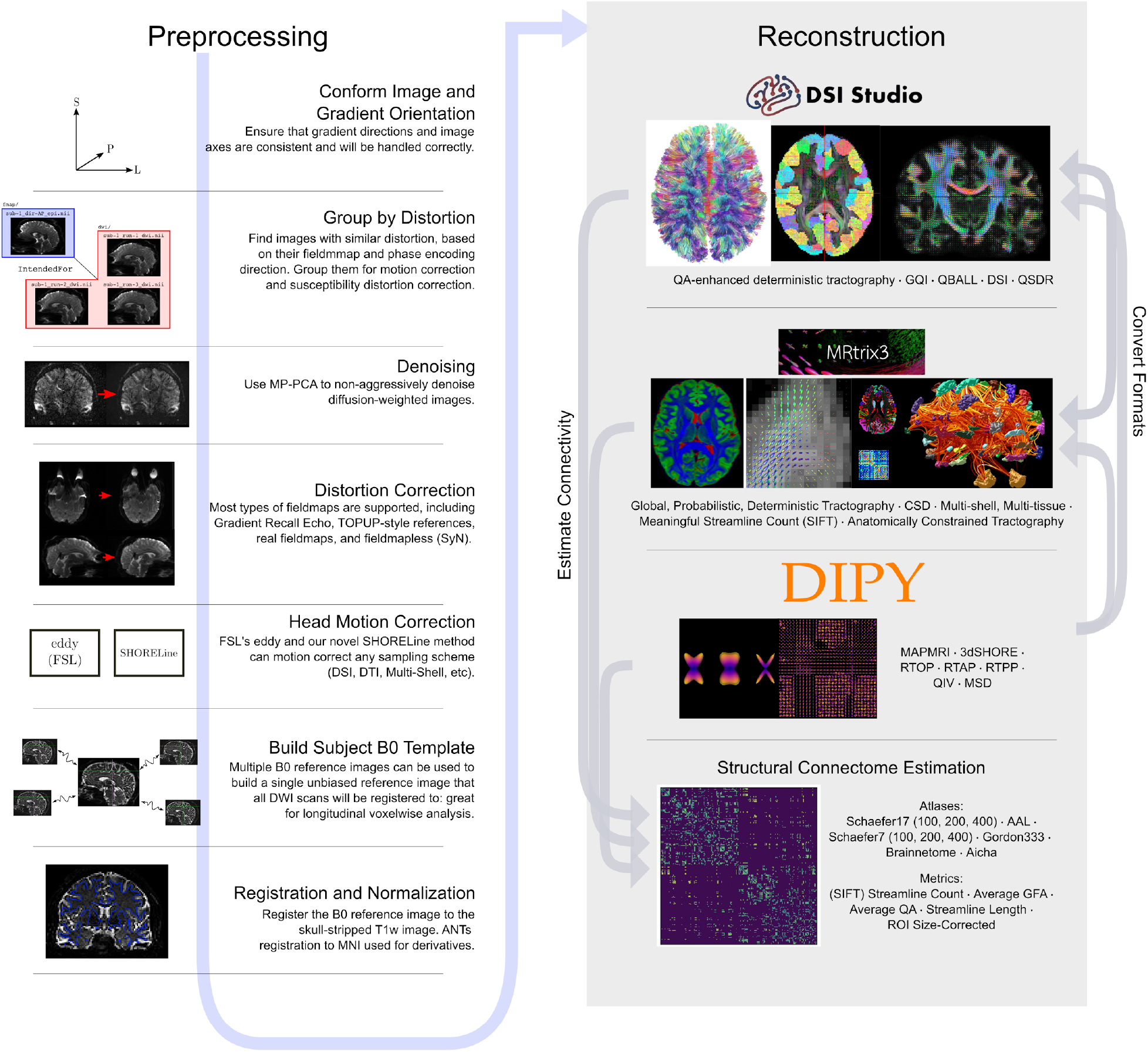
QSIPrep workflows. QSIPrep includes preprocessing (left column) and reconstruction (right column) workflows. BIDS data enters the workflow at the top left, following the blue arrow sequentially through the possible steps. The outputs from the preprocessing pipeline are inputs for the reconstruction workflows, which includes reconstruction methods from MRtrix3, DSI Studio, and DIPY. Gray arrows labeled “Estimate Connectivity” indicate that connectivity matrices can be estimated from all 3 software packages. Gray arrows labeled “Convert Formats” indicate that a reconstruction from one software package can be converted to be used in the destination software for further processing (e.g., DIPY reconstructions can be used for tractography in MRtrix3).

### QSIPrep provides intuitive visual reports and automated measures of data quality

The workflows used in QSIPrep automatically adapt to the input data specified in the BIDS layout provided by the input data. Therefore, it is critical for users to be able to see exactly which steps were taken and how these steps impact their data. To this end, QSIPrep produces HTML reports that describe how the input data were handled by the pipeline. Any step that alters the image data produces an animated “Before-After” visualization that allows users to visualize the transformation. Standardized text describing the workflow is automatically generated, including citations for any software or methods used; see **Supplementary Figure S1** and **Supplementary Note 2.1**. In addition, each preprocessed Diffusion-Weighted Image (DWI) series is accompanied by a quality control (QC) text file that describes the number of dropped slices, head motion summary measures, image dimensions before and after preprocessing, and a useful summary measure of data quality -- the neighboring DWI correlation (NDC)^16^.

Reconstruction workflows produce separate HTML reports that help the user quickly determine whether reconstruction was successful. Directional maxima are plotted in a mosaic of slices, along with Orientation Distribution Function (ODF) fields in a series of locations where different types of fiber crossings are visible^17^. If tractography was performed, the report includes an array of connectivity matrices, each using a different measure of connectivity (e.g., streamline count, length, and SIFT2 weights). A full example of a reconstruction report is provided in **Supplementary Figure S2;** an example of automatically generated methods text is provided in **Supplementary Note 2.2**.

### Preprocessing with QSIPrep reduces additional smoothing and enhances data quality

To demonstrate the generalizability of QSIPrep, we preprocessed nine different datasets acquired with a wide range of acquisition parameters and scanning platforms (n=657 total scans). These datasets included a standard single-shell sequence from the Philadelphia Neurodevelopmental Cohort (PNC)^18^, as well as four different multi-shell sampling schemes: the scheme used by the Adolescent Brain and Cognitive Development (ABCD) project^19^, a NODDI-optimized multi-shell acquisition^20^, The “Lifespan” sequence^21^ from the Human Connectome Project (HCP), and the multi-shell sequence collected by the Healthy Brain Network (HBN)^22^. Furthermore, we evaluated three Cartesian grid sampling schemes (DSI) with different sampling densities, as well as a compressed-sensing DSI (CS-DSI) sequence^23^ with random *q*-space sampling. A complete description of these datasets is presented in **Table 1**.

**Table 1.**
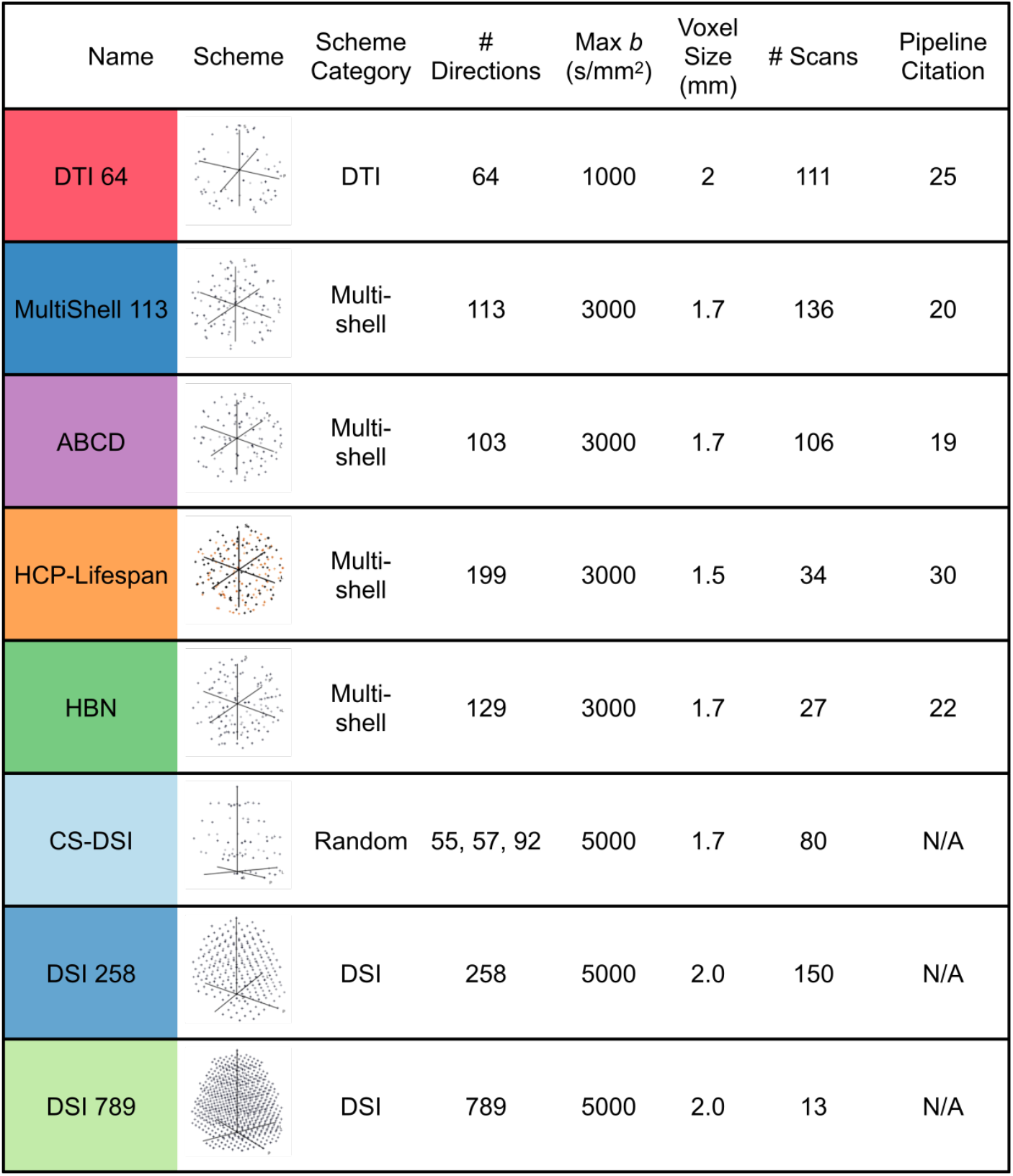
Diffusion imaging data used in QSIPrep development and evaluation. Cartesian (DSI), random (CS-DSI), and shelled (single-shell DTI and multi-shell) sequences were used to test the preprocessing and reconstruction workflows in QSIPrep. Sequences varied widely in their maximum gradient strength (1000-5000s/mm^2^), number of q-space samples (64-789) and voxel size (1.5-2.3mm). The row colors represent these schemes across all figures. The colors in the HCP-Lifespan image indicate that these samples came from different scans, grouped by phase-encoding direction.

| Name | Scheme | Scheme Category | # Directions | Max $b$ (s/mm<sup>2</sup>) | Voxel Size (mm) | # Scans | Pipeline Citation |
| --- | --- | --- | --- | --- | --- | --- | --- |
| DTI 64 |  | DTI | 64 | 1000 | 2 | 111 | 25 |
| MultiShell 113 |  | Multi-shell | 113 | 3000 | 1.7 | 136 | 20 |
| ABCD |  | Multi-shell | 103 | 3000 | 1.7 | 106 | 19 |
| HCP-Lifespan |  | Multi-shell | 199 | 3000 | 1.5 | 34 | 30 |
| HBN |  | Multi-shell | 129 | 3000 | 1.7 | 27 | 22 |
| CS-DSI |  | Random | 55, 57, 92 | 5000 | 1.7 | 80 | N/A |
| DSI 258 |  | DSI | 258 | 5000 | 2.0 | 150 | N/A |
| DSI 789 |  | DSI | 789 | 5000 | 2.0 | 13 | N/A |

For each of these datasets, we compared the performance of QSIPrep to that of published pipelines tailored for each dataset on two outcomes: image smoothness and image quality. The spatial smoothness of the *b*=0 images in each series* was characterized by calculating the mean of their estimated full width at half maximum (FWHM). This measure is impacted by multiple interpolations and imprecise spatial resampling of images, which introduce artifactual smoothness that reduces image contrast and anatomic detail. The quality metric we evaluated across datasets and pipelines was the NDC, a QC metric introduced in DSI Studio^10^.

Spatial smoothness and NDC scores provide complementary insights into how processing changes the raw images. NDC computes pairwise spatial correlation between each pair of dMRI volumes that sample the closest points in *q*-space. This computation can be applied to any *q*-space sampling scheme. Lower NDC values reflect reduced data quality, driven by noise and misalignment between dMRI volumes. While denoising and motion correction will increase NDC, it can also be artificially inflated by interpolation-driven spatial smoothing. Accordingly, we regressed image smoothness from the NDC values before comparing pipelines.

For shelled schemes, QSIPrep produced significantly less blurred images than pipelines tailored for many of the most widely used datasets (see **Figure 2a**, statistical results following Bonferroni correction in **Supplementary Table 1**). QSIPrep images were substantially less blurred than the custom pipelines developed for the single-shell DTI sequence from the PNC (*Δ*FWHM = −0.16mm), the multi-shell sequence from ABCD (*Δ*FWHM = −0.8mm), and the multi-shell sequence from the HCP-Lifespan (*Δ*FWHM = −0.75mm). Comparisons of raw and processed data further demonstrates the relatively large increase in smoothness introduced by many previously published pipelines (**Figure S4**). In contrast, the smoothness of QSIPrep’s outputs was slightly higher than that produced by the pipeline developed for the NODDI-optimized MultiShell 113 sequence (*Δ*FWHM = +0.09mm); no differences were seen in data from HBN.

**Fig. 2.**
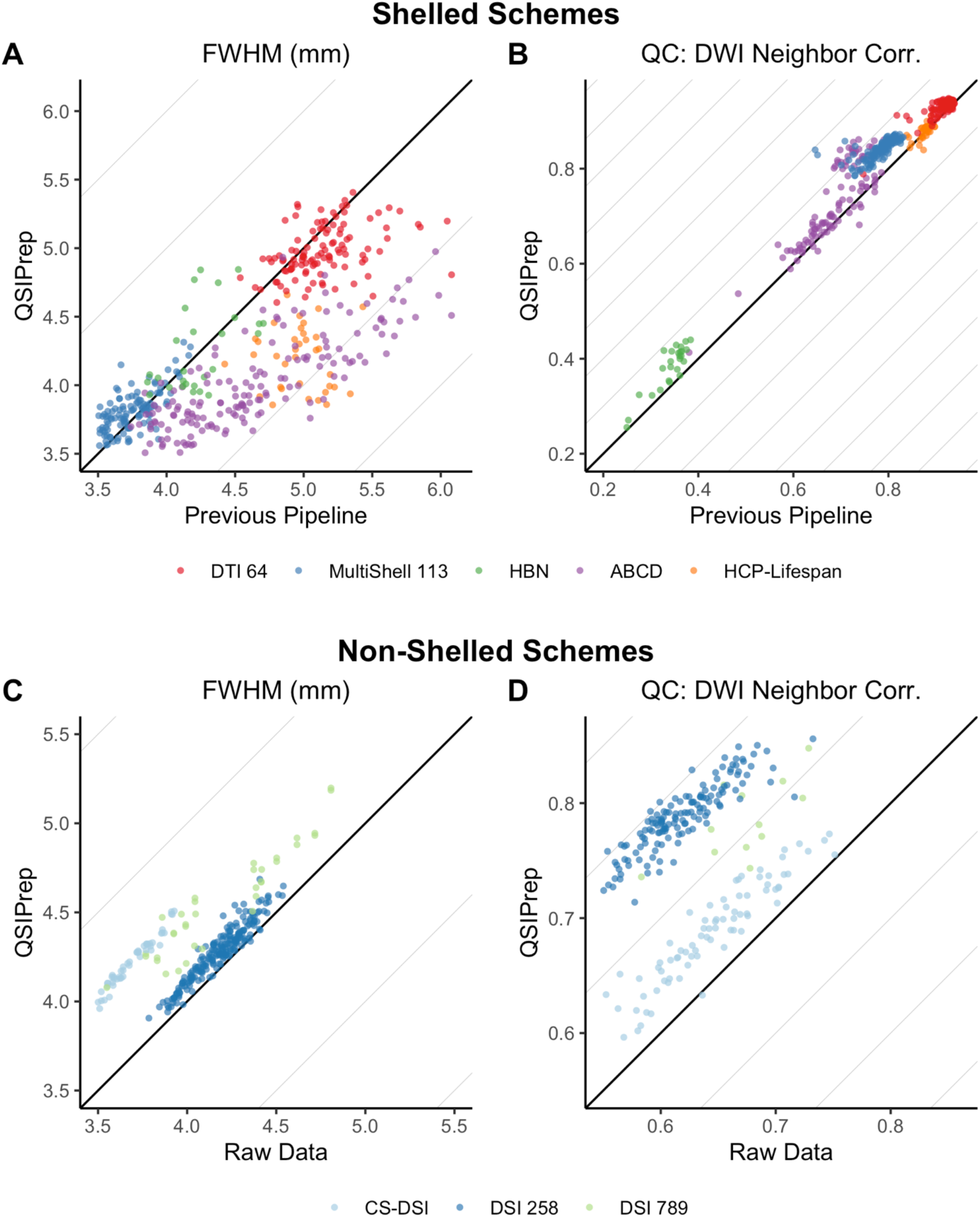
QSIPrep improves image quality without additional smoothing. For shelled schemes, image smoothness (FWHM) and data quality (neighboring DWI correlation; NDC) produced by QSIPrep were compared to previously published pipelines tailored for each acquisition scheme (see **Table 1**). QSIPrep introduced less blurring than published pipelines (panel **A**) in all cases except MultiShell 113 (where QSIPrep was slightly smoother) and HBN (where no difference was seen; see statistical results in **Supplementary Table 1**). Notably, QSIPrep yielded images with higher data quality than all pipelines evaluated, with the exception of the HCP-lifespan data, where no difference was present (panel **B**, see statistical results in **Supplementary Table 2**). For non-shelled schemes, as comparisons with existing pipelines were not available, QSIPrep was compared to raw data. As expected, compared to the raw data, non-shelled schemes were smoother after processing with QSIPrep (panel **C**; see statistical results in **Supplementary Table 3**). However, these differences were small (*Δ*FWHM < 0.5mm), especially when compared to the larger differences seen between existing pipelines developed for shelled schemes (i.e., *Δ*FWHM up to ~1.5mm in panel **A**; also see **Supplementary Figure 3** for analogous comparisons to raw data for shelled schemes). NDC QC scores were significantly improved for data processed with QSIPrep than the raw data for all non-shelled schemes (panel **D**; see statistical results in **Supplementary Table 4**).

Notably, QSIPrep yielded images with higher NDC than nearly all custom pipelines designed for shelled imaging sequences (see **Figure 2b** and statistical tests in **Supplementary Table 2**). The only exception to this was the HCP pipeline, where NDC scores were not significantly different from QSIPrep. These results emphasize that QSIPrep produces images of superior (or at least non-inferior) data quality compared to custom pipelines developed for a wide variety of shelled acquisition schemes.

One important advantage of QSIPrep is that in addition to shelled acquisition schemes, it can also effectively process advanced non-shelled schemes. In this case, no direct comparisons to an existing pipeline were available, so only comparisons with raw data were evaluated. Inevitably, any image processing introduces at least some increase in smoothness (see **Figure 2c** and statistical tests in **Supplementary Table 3**). As expected, images processed with QSIPrep were slightly but significantly smoother than the raw images (*Δ*FWHM = +0.53mm), similar to that seen for shelled schemes (see **Supplementary Figure 3**). Notably, processing non-shelled sequences with QSIPrep significantly improved data quality, reflected in a large increase in NDC values (see **Figure 2d** and statistical tests in **Supplementary Table 4**).

### Interoperability of reconstruction workflows enables direct comparison of disparate methods

A major challenge in comparing reconstruction methods is that many dMRI software packages have their own file formats, coordinate systems, orientation conventions, and visualization tools (see **Supplementary Note 1**). This diversity is compounded by the large number of possible dMRI acquisition schemes, many of which only meet the requirements of a subset of reconstruction methods. QSIPrep’s set of curated reconstruction workflows provides two critical benefits to users: correct processing and a uniform derived output format. First, workflows are designed to ensure that data preprocessed by QSIPrep are handled correctly *within* the reconstruction workflow. Second, outputs from each reconstruction method conform to a consistent format *across* methods.

This emphasis on software interoperability facilitates comparisons between methods. For example, **Figure 3** displays the results from a number of reconstruction workflows, depicting disparate sampling schemes reconstructed using popular methods from MRtrix3, DSI Studio, and the Laplacian-regularized MAPMRI (MAPL)^24^ implementation from DIPY. The visual similarity of the reconstructed ODFs and Fiber Orientation Distributions (FODs) suggests that many of these methods share important features like peak directions. All reconstruction outputs are produced in the native file format of each software package used and also consistently provided in a DSI Studio (fib format) file.

**Fig 3.**
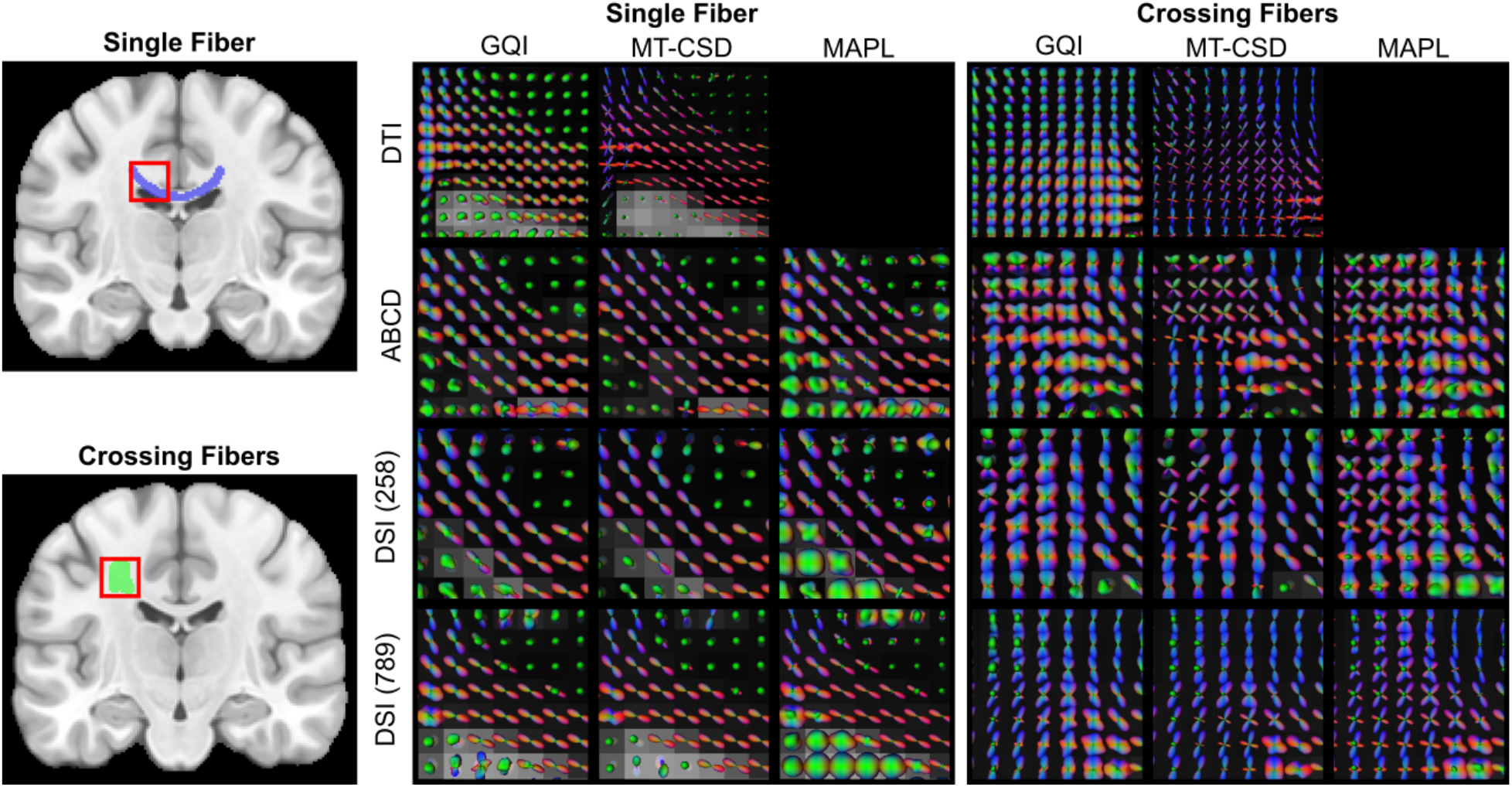
QSIPrep reconstruction workflows produce comparable output across diverse sampling schemes and reconstruction methods. Four sampling schemes each reconstructed using four methods: GQI from DSI Studio, multi-tissue CSD from MRtrix, and MAPL from Dipy. ODF fields are shown in two white matter regions (left), a single fiber area in the corpus callosum (top) and a crossing fiber region in the centrum semiovale (bottom). The middle panel shows ODFs reconstructed in the single fiber region, and the right panel shows ODFs reconstructed in the crossing fiber region for the four sampling schemes (rows) and the three reconstruction methods (columns). The ability to run multi-tissue CSD on DSI data is a unique feature of QSIPrep.

Capitalizing upon the interoperability described above, QSIPrep also allows users to apply standard processing and reconstruction methods developed for shelled sequences to advanced non-shelled sequences. To do this, QSIPrep converts non-shelled sampling schemes to a multi-shell scheme using a 3dSHORE-based *q*-space interpolation. This conversion allows, for example, the use of multi-shell multi-tissue reconstruction and MRtrix3 tractography methods on *any* non-shelled sampling scheme. The ability to apply standard analytic methods to non-shelled schemes dramatically increases the accessibility of these advanced acquisition sequences.

### Structural connectome estimation

One of the most popular applications for dMRI is to construct whole-brain structural connectomes via streamline tractography. However, file formats for storing and representing connectomes vary across software packages, thereby limiting comparisons. For example, MRtrix3 produces text files, DSI Studio produces MATLAB files, and DIPY produces NumPy arrays. Furthermore, many software packages produce inconsistently sized matrices across subjects, due to some participants missing small regions from high-resolution atlases. In contrast, QSIPrep ensures that connectivity matrices are directly comparable across methods and participants. Specifically, the software checks that matrices are correctly shaped across all atlases and stores them in easily accessible HDF5 files. Finally, due to the interoperability of the component software elements, QSIPrep allows a far more diverse array of connectivity measurements to be calculated than is possible with individual software packages (see **Table 2**).

**Table 2.** Reconstruction workflows included with QSIPrep. Supported reconstruction workflows in QSIPrep version 0.9.0. Workflow names indicate the main software package used for the workflow. DSI Studio *connectivity Weights* include for each region pair: streamline count, length-normalized streamline count, average along-streamline GFA and mean streamline length. MRtrix3 *connectivity weights* include SIFT2-weighted streamline counts (as well as the inverse node-volume corrected version), streamline count, and mean streamline length.

| Name | Supported Schemes | Scalar maps | Tractography | Connectivity weights |
| --- | --- | --- | --- | --- |
| mrtrix_multishell_msmt | Multi-shell | None | Probabilistic | MRtrix3 |
| mrtrix_singleshell_ss3t | Single-shell | None | Probabilistic | MRtrix3 |
| dsi_studio_gqi | Multi-shell<br>Single-shell<br>Cartesian | ISO<br>GFA<br>QA[0-3] | Deterministic | DSI Studio |
| dipy_mapmri | Multi-shell<br>Cartesian<br>Random | RTOP<br>RTAP<br>RTPP<br>MSD QIV | Both | DSI Studio |
| dipy_3dshore | Multi-shell<br>Cartesian<br>Random | RTOP<br>SHORE<br>Coefs | Both | DSI Studio |
| cdsdi_3dshore | Cartesian<br>Random | None | Both | MRtrix3<br>DSI Studio |

## DISCUSSION

QSIPrep provides a fully reproducible and transparent platform for preprocessing and reconstructing virtually all dMRI data. It is unique in that it integrates cutting-edge methods from multiple software packages to build proper preprocessing and reconstruction pipelines for all nearly all diffusion sampling schemes. The use of complementary tools incorporated by QSIPrep likely contributes to our finding that its automatically configured pipelines perform as well or better than published workflows tailored to each dataset.

The generalizability of QSIPrep is facilitated by two major features. As a BIDS app, the processing workflow adapts to the characteristics of the input data, producing an appropriate pipeline as long as the user has correctly specified their data in BIDS. The BIDS app interface alleviates much of the burden for users who wish to follow best practices in data processing, but do not have the time or skills to learn the minutiae of multiple software packages. The adaptive pipelines configured by QSIPrep dramatically enhances accessibility and reproducibility without sacrificing quality. This is underscored by the result that pipelines automatically constructed by QSIPrep yielded results with comparable or better data quality and smoothness when compared to established pipelines for multiple studies.

Notably, QSIPrep is distributed as both a Python package and as a Docker container that includes all the dependencies, ensuring that it is able to run on all modern computing systems. QSIPrep’s development is open and uses GitHub to manage feature requests, bug reports, and user questions. Basic functionality is constantly verified using continuous integration testing, yielding containers whose contents are curated by numbered releases that chronicle bug fixes and new features. This design philosophy and development practice has enabled QSIPrep to rapidly respond to bugs and feature requests, accelerating the growth of its user and developer base.

While dMRI presents unique challenges, such as diversity of sampling schemes and software packages, it also has advantages that allow for scalability. One major obstacle to scalability in large-scale imaging efforts is consistent, quantitative measures of image quality. Notably, QSIPrep calculates the NDC, in-scanner motion, the number of dropped slices, and the *b*=0 intensity variability^25^ as quality metrics. All measures are scalar values that can be used to quickly assess the relative quality of scans regardless of sampling scheme, so outlying values can be flagged for detailed examination of the rich HTML reports that QSIPrep generates.

QSIPrep offers a particularly large advance in processing cutting-edge Cartesian and random *q*-space sampling schemes, which previously had no publicly available head motion correction methods. Cartesian sampling schemes are unique in providing a direct relationship to the diffusion propagator^2^ and random sampling schemes inherit this advantage while taking much less time to acquire^1^. QSIPrep provides novel methods for preprocessing these valuable images and allows users to apply tractography and reconstruction methods previously only available to shelled sampling schemes. These advances enhance the accessibility of advanced sampling schemes to the neuroscience community.

Several limitations of the current version of QSIPrep should be noted. First, the software does not support double diffusion encoding *q*-space imaging or gradient tensor imaging. These scanning sequences are not widely used, not currently supported by BIDS, and lack open preprocessing software and methods. Second, it is critical to note that we do not claim that the reconstruction workflows are optimal for any given method, only that they implement current best practices. Traditionally, the question of optimality in reconstruction and tractography methods has been difficult to address, in part due to the lack of comparability of measures produced by different software packages. The interoperability provided by QSIPrep facilitates the comparison of many measures – including ODFs, anisotropy scalars, and connectivity matrices – across reconstruction methods and sampling schemes.

Taken together, QSIPrep allows researchers to correctly apply reproducible preprocessing pipelines and advanced reconstruction methods to nearly any dMRI data. By harnessing cutting-edge techniques from individual software packages and unifying them in an interoperable framework, the widely generalizable methods provided by QSIPrep perform as well or better than existing customized solutions that can only be applied to a subset of sampling schemes. As such, QSIPrep facilitates the adoption of fully reproducible best practices for the processing, quality assurance, and reconstruction of diffusion images.

## ONLINE METHODS

### The QSIPrep workflow

The preprocessing workflow is dynamically built based on data provided as BIDS input. Separate dMRI scans can be grouped and processed together depending on their acquisition parameters and user-supplied options. Image processing can include denoising, head motion, eddy current and distortion correction, b=0 reference image creation (including an optional single-subject b=0 template), coregistration to the T1w image, spatial normalization, image resampling, and gradient rotation. **Figure 1**’s left panel depicts the sequence of these steps.

#### Conform, Merge, and Denoise workflow

One of the unique challenges of dMRI preprocessing is that the *q*-space sampling scheme is often split into multiple separate scans. Moreover, groups of these scans may be acquired with opposite phase encoding directions so that their *b*=0 images can be used for SDC. The heuristic used by QSIPrep is to divide the scans into “warped groups” that share the same susceptibility distortions. The warped groups are sent to the conform, merge, and denoise workflow.

All spatial transformation operations in QSIPrep (excluding TOPUP/eddy) are performed using ANTs^15^. ANTs internally uses an LPS+ coordinate system. The FSL-style bvec format required by BIDS specifies gradient directions with respect to the image axis, not world coordinates. By conforming all images and bvecs to LPS+ image orientation, ANTs can be used directly for registration and transformation on both the images and the gradient vectors. The *conform* step enforces this orientation and checks that the images have matching qform/sform mappings.

Next, warped groups are denoised (using MP-PCA^26^, Gibbs unringing^27^, bias correction^28^, *b*=0 image-based intensity normalization), and concatenated if multiple runs are present. This step can be done as concatenate-then-denoise or denoise-then-concatenate (default), depending on the user’s preference. If images are concatenated before denoising, there will be more data for MP-PCA to include in its denoising. However, if the concatenated scans are very far out of alignment with one another, the performance of MP-PCA may be sub-optimal. The other denoising methods are not affected by when data is concatenated. The user can select the concatenate-then-denoise order using a command line flag. A visual description of these workflows is presented in **Supplementary Figure 3**.

#### HMC/ECC/SDC workflow

We combined head motion correction, eddy current correction, and susceptibility distortion into a single workflow due to the interdependence of the TOPUP and eddy tools. This workflow is split into special cases for shelled sampling schemes (multi-shell or single-shell) and all other sampling schemes.

##### Shelled sampling schemes

If a reverse-phase encoding direction image is available in the `fmap`/ or `dwi`/ directories, a fieldmap is calculated using TOPUP and sent to eddy to be applied in addition to HMC and ECC. In all other cases the fieldmap is calculated using workflows adapted from fMRIPrep and applied to the motion-corrected output from eddy.

##### Cartesian and random sampling schemes

These schemes are processed using the QSIPrep’s novel SHORELine algorithm^†^ before being processed using the distortion correction workflows.

Regardless of the sampling schemes, SDC requires a careful selection of representative *b*=0 images from each DWI scan. QSIPrep selects up to three (depending on availability) *b*=0 images evenly spaced in time from each group of phase encoding directions. Using a representative subset of all *b*=0 images is required to limit the run time of TOPUP. The details of which images are used for SDC are included in the HTML report.

#### b=0 template workflow

The reference image for each DWI series is created by extracting the *b*=0 images from the series after HMC, ECC, and SDC. They are combined using a normalized average as implemented in ANTs and undergo a histogram equalization as implemented in DIPY. A visual report is generated showing the *b*=0 template before and after histogram equalization.

#### Intramodal template workflow

In cases where there are multiple sessions or multiple separate DWI scans that should not be merged, there will be multiple *b*=0 reference images. Each can be affected by errors in SDC or intermodal coregistration to the T1w image. QSIPrep provides the option to create an “intramodal template” using ANTs template construction^29^ on the set of *b*=0 reference images. The intramodal template is co-registered to the T1w image instead of each individual *b*=0 reference image. The transform to the intramodal template and as well as the intramodal template’s transform to register to the T1w image are added to the stack of transforms that get combined and applied to each DWI.

#### Coregistration and resampling workflow

Coregistration between the *b*=0 template images (or the intramodal *b*=0 template) is performed using `antsRegistration`. If the user requests a T1w-based spatial normalization to a template, this is also performed using the `antsRegistration`-based workflow from fMRIPrep. Similar to the HCP Pipelines^30^ and the ABCD MMPS pipeline^19^, QSIPrep uses a rigid transformation to register the skull-stripped T1w image to AC-PC alignment. Unlike these other pipelines, QSIPrep combines all spatial transformations so that only a single resampling is ever performed on the images. The final resampling uses a Lanczos-windowed Sinc interpolation if the requested output resolution is close to the resolution of the input data. If more than a 10% increase in spatial resolution is requested, then a BSpline interpolation is performed (as suggested in the MRtrix3 documentation). The final resampling can at most include the affine head motion correction, the polynomial eddy current correction, the nonlinear susceptibility distortion correction, the nonlinear registration to the *b*=0 template, the coregistration to the T1w image, and the realignment to AC/PC orientation. Combining these into a single shot interpolation helps preserve high frequency spatial features.

### QSIPrep reconstruction workflows

QSIPrep’s curated reconstruction workflows apply developer-recommended post-processing and reconstruction steps, storing the results in both the software-native and DSI Studio formats. The pipelines were chosen from the most popular open source diffusion imaging software packages such that there is at least one workflow for each *q*-space sampling scheme. A comparison of pipelines is shown in **Table 2** and their implementation details are described below grouped by software. While the included workflows use fixed parameters, users can download and edit workflow configuration files to change the workflow’s behavior.

#### MRtrix3

There are a number of MRtrix3-based workflows that share the same initial steps but differ in how the FOD estimation is performed. In each MRtrix3-based workflow the fiber response function is estimated using `dwi2response dhollander`^31^ with a brain mask based on the T1w. The main differences are the MRtrix3 workflows are in 1) the CSD algorithm used to estimate WM FODs and GM/CSF compartments (either multi-shell multi-tissue CSD, MSMT-CSD; or single-shell 3-tissue^32,33^ CSD, SS3T-CSD) and 2) whether a T1w-based tissue segmentation is used during tractography. In the `*_noACT` versions of the pipelines no T1w-based segmentation is used during tractography, which is desirable if no SDC was performed during preprocessing. Otherwise, cropping is performed at the GM/WM interface along with backtracking. In all MRtrix3 pipelines, tractography is performed using tckgen, which employs the iFOD2 probabilistic tracking method to generate 10^7^ streamlines with a maximum length of 250mm, minimum length of 30mm, FOD power of 0.33. Weights for each streamline are calculated using SIFT2, which is then used to estimate the structural connectivity matrix.

##### mrtrix_multishell_msmt

This workflow uses the `dwi2fod msmt_csd` algorithm^31^ to estimate FODs for white matter, gray matter and cerebrospinal fluid using multi-shell acquisitions. The white matter FODs are used for tractography and the T1w segmentation is used for anatomical constraints^34^. *mrtrix_multishell_msmt_noACT* is identical except that no T1w-based anatomical constraints are used in tractography. *mrtrix_singleshell_ss3t* is optimized for single-shell acquisitions and also estimates multi-tissue FODs for white matter, gray matter and cerebrospinal fluid using the `ss3t_csd_beta1` (SS3T-CSD) algorithm^32,33^, provided via the MRtrix3Tissue fork of MRtrix3. The white matter FODs are used for tractography and the T1w segmentation is used for anatomical constraints^34^. *mrtrix_singleshell_ss3t_noACT* removes the anatomical constraints from tractography.

#### DSI Studio

*dsi_studio_gqi* runs the standard GQI reconstruction^35^ followed by deterministic tractography^36^. GQI works on almost any sampling scheme. GQI models the diffusion ODF, so ODF peaks are much smaller than is commonly seen with CSD. Diffusion ODFs exhibit relatively small ODF peaks, yet still robustly detect fiber crossings^37^. Although GQI technically works on DTI scans, with spherical sampling on a single-shell around *b*=1000 s/mm^2^, its performance markedly improves when more *q*-space samples are available. The tractography performed in this pipeline ensures that 5 million streamlines are created with a maximum length of 250mm, a minimum length of 30mm, random seeding, a step size of 1mm, and an automatically calculated QA threshold. Additionally, a number of anisotropy scalar images are produced such as quantitative anisotropy (QA)^35^, generalized fractional anisotropy (GFA), and the isotropic component of the ODF.

#### DIPY

*dipy_mapmri* Mean Apparent Propagator MRI (MAPMRI) is a recently proposed reconstruction method^4^ that can estimate ensemble average diffusion propagators (EAPs) and ODFs analytically using multi-shell, Cartesian or random *q*-space sampling schemes. This method produces EAP-derived scalars like return to origin probability (RTOP), return to axis probability (RTAP), return to plane probability (RTPP), *q*-space inverse variance (QIV), and mean squared displacement (MSD). The ODFs are saved in DSI Studio format and optionally as spherical harmonics coefficients in the MRtrix3 format. *dipy_3dshore.* The 3D Simple Harmonic Oscillator-based Reconstruction and Estimation (3dSHORE)^38^ method also uses a closed-form solution to estimate EAPs and ODFs from *q*-space data. This workflow uses the BrainSuite 3dSHORE basis in a DIPY reconstruction. Much like *dipy_mapmri*, EAP-related scalars such as RTOP, RTAP, RTPP, and MSD are estimated. For both of these reconstruction pipelines, tractography is run identically to the *dsi_studio_gqi*.

#### Experimental DSI scheme-converting reconstruction

*`csdsi_3dshore`* This pipeline is for DSI or compressed-sensing DSI. The first step is an L2-regularized 3dSHORE reconstruction^24^ of the ensemble average propagator in each voxel. These EAPs are then used to 1) calculate ODFs, which are then sent to DSI Studio for tractography and 2) impute signal for a multi-shell (specifically HCP) sampling scheme, which is run through the `mrtrix_multishell_msmt` pipeline. The resampling is similar to a previously-described GQI-based method^39^ but uses the 3dSHORE basis set to estimate out-of-sample images.

#### Structural connectivity matrices

Tractography resulting in connectivity matrices are conformed to a standard HDF5-based output format so as to be directly comparable across methods and software packages. A set of commonly-used parcellation schemes are included with QSIPrep, such as the Schaefer atlases in the 100, 200, and 400 resolutions, the brainnetome atlas (264 regions), AICHA (384 regions), Gordon (333 regions), the AAL (116 regions), and the Power atlas (264 regions). Furthermore, users can easily add their own custom atlases as required.

### Evaluation Data

Data were gathered from a number of independent studies from multiple institutions. These samples were selected to test a variety of *q*-space sampling schemes and evaluate if QSIPrep handles each one correctly. An overview of the acquisition parameters is provided in **Table 1**. QSIPrep was run on the raw data from each study. Spatial smoothness and neighboring DWI correlation^17^ were calculated for the QSIPrep-preprocessed data and for the data processed using a pipeline specifically designed for that sampling scheme. In the case of non-shelled schemes, QSIPrep was compared to unprocessed data.

#### Single-shell DTI

The single shell data was collected as part of the Philadelphia Neurodevelopmental Cohort (PNC)^18^ and processed according to the methods described by Roalf *et al.*^25^. This pipeline is similar to QSIPrep, utilizing eddy (from FSL5) and custom code for applying distortion correction. The QSIPrep pipeline differs in that it adds MP-PCA, Gibbs unringing, FSL6, and resampling using ANTs. A total of 111 subjects were randomly selected from the available PNC dMRI data.

#### Multi-shell, NODDI-optimized

This sampling scheme was designed with the goal of fitting microstructural models such as NODDI^5^. The data were published Pines *et al*.^20^, where the preprocessing scheme used FSL5’s TOPUP and eddy with outlier replacement enabled. The QSIPrep pipeline differed in that it added MP-PCA denoising, Gibbs unringing, FSL6, and resampling using ANTs. We evaluated a sample of 136 participants from the cohort used by Pines *et al.*.

#### Multi-shell, ABCD

A total of 106 datasets were downloaded from the NIMH Data Archive (NDA) repository in their converted-to-NIfTI and minimally preprocessed form. The ABCD dMRI preprocessing pipeline^19^ does not use any of the same software as QSIPrep but performs similar steps. The ABCD pipeline includes gradient nonlinearity correction and uses in-house code for performing Eddy current and distortion correction. QSIPrep adds MP-PCA, Gibbs unringing, ECC, and SDC using FSL6 and resampling using ANTs.

#### Multi-shell HCP-Lifespan

A total of 34 subjects were scanned using the HCP-Lifespan imaging protocol^21^ and processed using both the official HCP diffusion pipelines^30^ (v4.0.0-alpha.5) and QSIPrep. The HCP diffusion pipeline included motion and eddy current correction, distortion correction, across-scan intensity normalization, coregistration to the T1w image, gradient unwarping and image pair averaging. QSIPrep was upgraded as part of 0.9.0beta1 to include the image pair averaging so that QC measures could be compared directly between the QSIPrep and HCP pipeline outputs. QSIPrep was adjusted to use a quadratic first-level model in *eddy* to match the HCP diffusion pipeline.

#### Multi-shell HBN

A total of 27 HBN^22^ subjects were processed using both an early prototype version of dMRIPrep (https://github.com/nipy/dmriprep) and QSIPrep. Both dMRIPrep and QSIPrep use TOPUP and eddy for distortion, eddy current, and motion correction, but dMRIPrep did not include Gibbs unringing or MP-PCA.

#### Cartesian grid (DSI) schemes

Prior to QSIPrep there was no publicly available software for applying head motion correction to DSI or CS-DSI acquisitions. Therefore, the QSIPrep-preprocessed images were compared directly to the NDC calculated on the raw images.

### Evaluation Data Analysis

#### Smoothness comparison

Smoothness was estimated using AFNI’s `3dFWHMx` on each b=0 image from the QSIPrep-preprocessed images and the preprocessed images from the previously used pipelines. A linear mixed-effects model was used to evaluate the impact of pipeline on the outcome of image smoothness (FWHM) with subject as a random intercept. Models were fit separately for each sampling scheme; Bonferroni correction was used to correct for multiple comparisons. For shelled schemes, comparisons were made versus a previously published existing pipeline tailored for that sequence. For non-shelled images, due to the lack of well-documented existing pipelines, QSIPrep was compared to the raw images.

#### Neighboring DWI Correlation (NDC) comparison

NDC can also be artificially inflated by interpolation-driven spatial smoothing. Accordingly, we regressed image smoothness (FWHM) from the NDC values before comparing pipelines. The FWHM-corrected NDC was modeled as a function of pipeline and subject, where the pipeline was a factor, and the subject was a random intercept. As for smoothness, models were fit separately for each dataset, and multiple comparisons were corrected using the Bonferroni method.

#### Code availability

All data and code used to perform these tests are available at: https://pennlinc.github.io/qsiprep_paper/ (DOI: 10.5281/zenodo.4014341)

## Footnotes

* We did not include *b*>0 images because their spatial frequencies strongly depend on gradient strength and direction. The *b*=0 images are directly comparable across all sampling schemes.

† https://qsiprep.readthedocs.io/en/latest/preprocessing.html?#head-motion-estimation-shoreline

## Notes

### Competing Interest Statement

The authors have declared no competing interest.

