## Supplementary material for "QSIPrep: An integrative platform for preprocessing and reconstructing diffusion MRI": Online Supplementary Materials

| Scheme | Estimate | Std. Error | t-Statistic | p-Value (Bonf.) |
|---|---|---|---|---|
| DTI 64 | -0.163 | 0.025 | -6.47 | < 0.001 |
| ABCD | -0.794 | 0.03 | -26.284 | < 0.001 |
| HBN | 0.018 | 0.053 | 0.345 | 0.733 |
| HCP-Lifespan | -0.751 | 0.053 | -14.089 | < 0.001 |
| MultiShell 113 | 0.092 | 0.012 | 7.639 | < 0.001 |

**Table S1 | FWHM Comparison: Shelled.** Negative values in "Estimate" indicate that QSIPrep produced sharper (less-smooth) images than the previous pipeline. Linear mixed models revealed that QSIPrep was sharper than all other pipelines except for MultiShell 113 (which was less smooth by 0.092mm) and HBN, where there was no significant difference.

| Scheme | Estimate | Std. Error | t-Statistic | p-Value (Bonf.) |
|---|---|---|---|---|
| DTI 64 | 0.015 | 0.001 | 9.984 | < 0.001 |
| ABCD | 0.041 | 0.004 | 9.714 | < 0.001 |
| HBN | 0.041 | 0.004 | 9.341 | < 0.001 |
| HCP-Lifespan | 0.002 | 0.002 | 1.097 | 0.281 |
| MultiShell 113 | 0.058 | 0.002 | 30.101 | < 0.001 |

**Table S2 | QC Comparison: Shelled.** Positive values in “Estimate” reflect higher FWHM-corrected NDC QC values in QSIPrep. Linear mixed effects models revealed that QSIPrep produced significantly higher data quality than all pipelines other than the HCP Diffusion pipeline, where there was no significant difference.

| Scheme | Estimate | Std. Error | t-Statistic | p-Value (Bonf.) |
|---|---|---|---|---|
| CS-DSI | 0.528 | 0.004 | 140.789 | < 0.001 |
| DSI 258 | 0.099 | 0.004 | 22.716 | < 0.001 |
| DSI 789 | 0.368 | 0.03 | 12.159 | < 0.001 |

**Table S3 | FWHM Comparison: Non-Shelled.** Here "Estimate" is the increase in FWHM (in mm) from the raw images to the output of QSIPrep. Linear mixed models revealed that preprocessed images are statistically significantly smoother than the raw data, but the difference was small (at most 0.53mm).

| Scheme | Estimate | Std. Error | t-Statistic | p-Value (Bonf.) |
|---|---|---|---|---|
| CS-DSI | 0.043 | 0.002 | 22.275 | < 0.001 |
| DSI 258 | 0.17 | 0.001 | 120.015 | < 0.001 |
| DSI 789 | 0.113 | 0.008 | 13.99 | < 0.001 |

**Table S4 | QC Comparison: Non-Shelled.** Positive values in "Estimate" indicate QSIPrep had higher FWHM-corrected NDC QC scores than the raw data. Linear mixed models revealed that QSIPrep produced significantly larger values in all cases.

Actual report from QSIPrep

Description of report content

### Anatomical

#### Anatomical Conformation

- Input T1w images: 1
- Output orientation: LPS
- Output dimensions: 192x256x160
- Output voxel size: 0.94mm x 0.94mm x 1mm
- Discarded images: 0

#### Brain mask and brain tissue segmentation of the T1w

This panel shows the template T1-weighted image (if several T1w images were found), with contours delineating the detected brain mask and brain tissue segmentations.

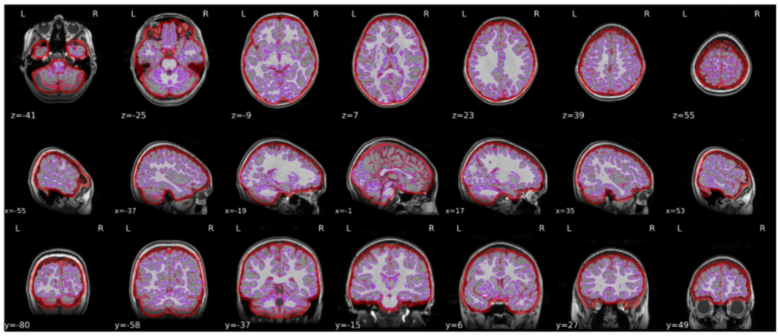

Get figure file: sub-113492/figures/sub-113492_seg_brainmask.svg

Anatomical preprocessing and reports are adopted from fMRIPrep.

#### T1 to MNI registration

Nonlinear mapping of the T1w image into MNI space. Hover on the panel with the mouse to transition between both spaces.

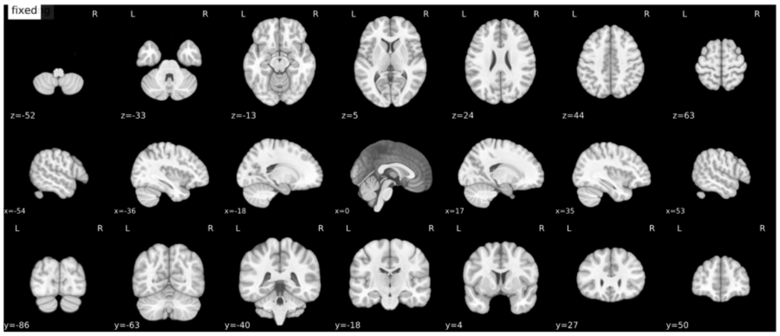


Get figure file: sub-113492/figures/sub-113492_t1_2_mni.svg

This spatial normalization is to be used only for mapping atlases from template space to the preprocessed dMRI data.

Spatially normalizing dMRI data should occur after reconstruction.

### Denoising

#### Reports for Session: PNC1 Run: 01

##### MP-PCA denoising

Effect of MP-PCA denoising on a low and high-b image.

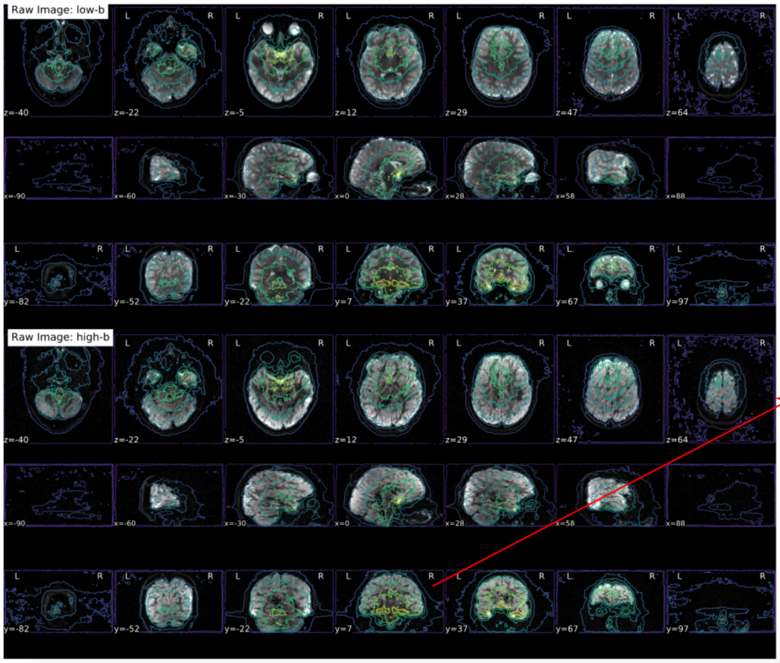


Images are shown before and after MP-PCA denoising. A contour plot of the noise level can be used to check if noise levels correspond to anatomical features.

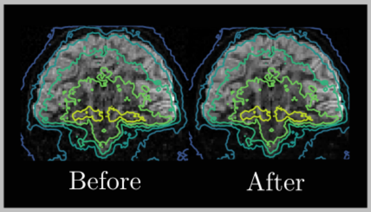


##### Gibbs Ringing Removal

Effect of removing Gibbs ringing on a low and high-b image.

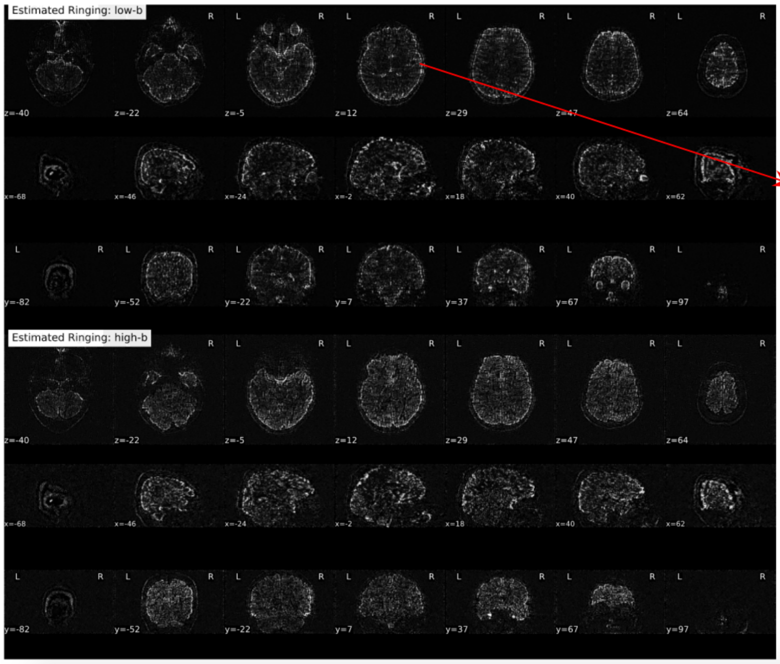


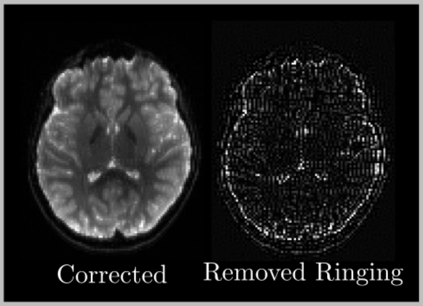


The effects of running the unringing algorithm implemented in MRtrix3's mrdegibbs program is shown. The animation flashes between the corrected image (above left) and the estimated artifact that was subtracted (above right).

##### DWI Bias correction

Effect of bias correction on a low and high-b image. Bias field contour lines are drawn as an overlay.

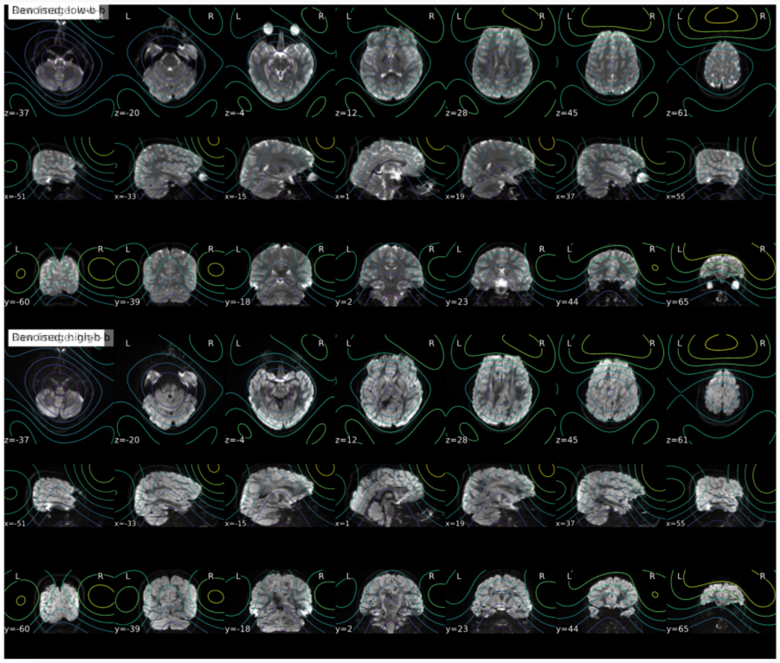

##### b=0 Reference Image

b=0 template and final mask output. The t1 and signal intersection mask is blue, their xor is red and the entire mask is plotted in cyan.

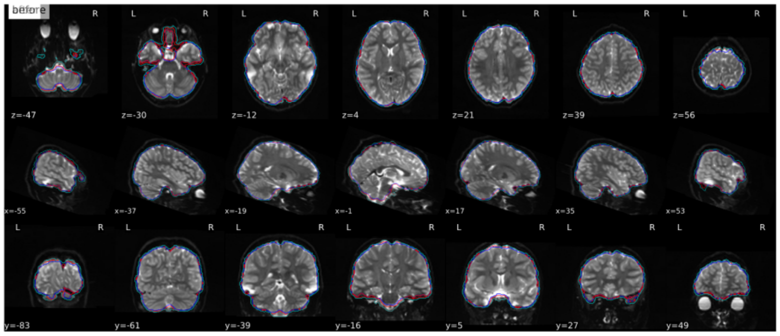

Spatial bias correction is performed using MRtrix3's dwibiascorr. This report shows a mosaic of before and after the correction is applied.

Contour lines of the spatial bias field are shown over the images.

The b=0 reference is critical for coregistration and masking. It is estimated based on the motion-corrected b=0 images in the DWI series.

QSIPrep masks the b=0 reference based on its signal and the mask based on the T1-weighted image.

#### Fieldmap

Overlaid on the reference EPI image

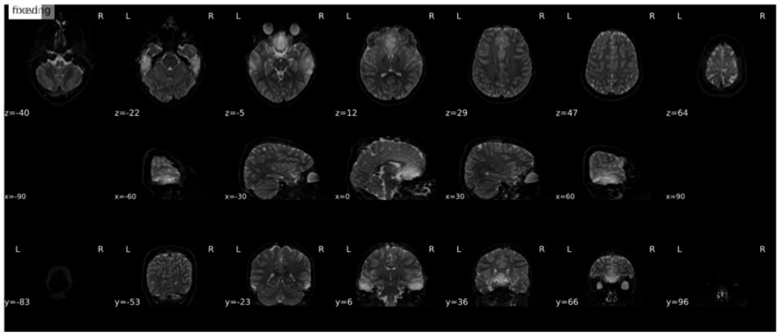

Fieldmaps are calculated using workflows from fMRIPrep. Here the fieldmap in Hz is shown on the b=0 reference image.

#### Susceptibility distortion correction

Results of performing susceptibility distortion correction (SDC) on the EPI

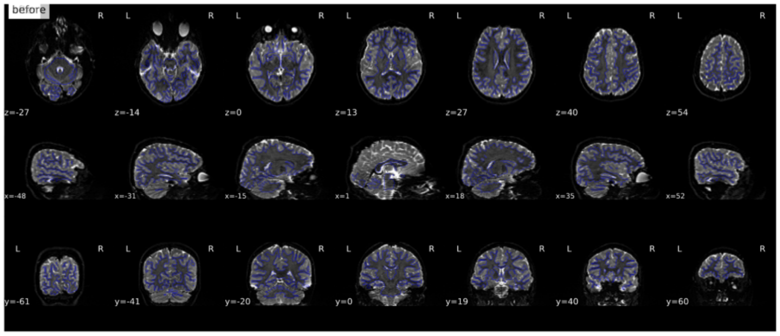

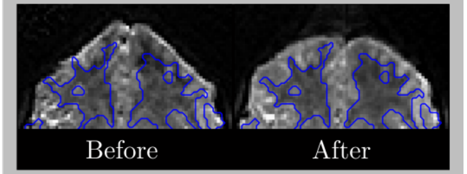


An animation of the b=0 reference image before and after applying SDC. The gray/white matter interface is drawn as a blue contour line.

#### b0 to T1 registration

`antsRegistration` was used to generate transformations from b0I-space to T1w-space

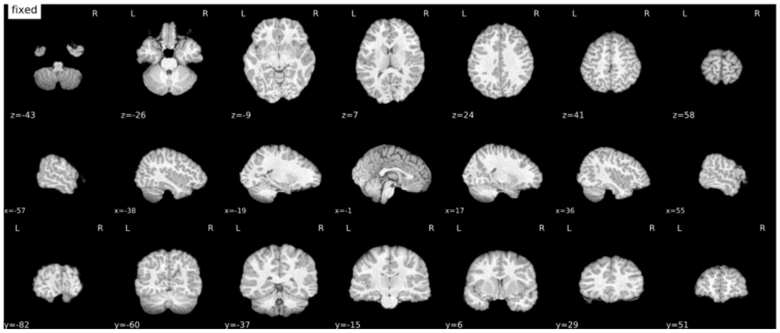

Intermodal registration is performed using ANTs. This animation shows the b=0 template relative to the T1w brain.

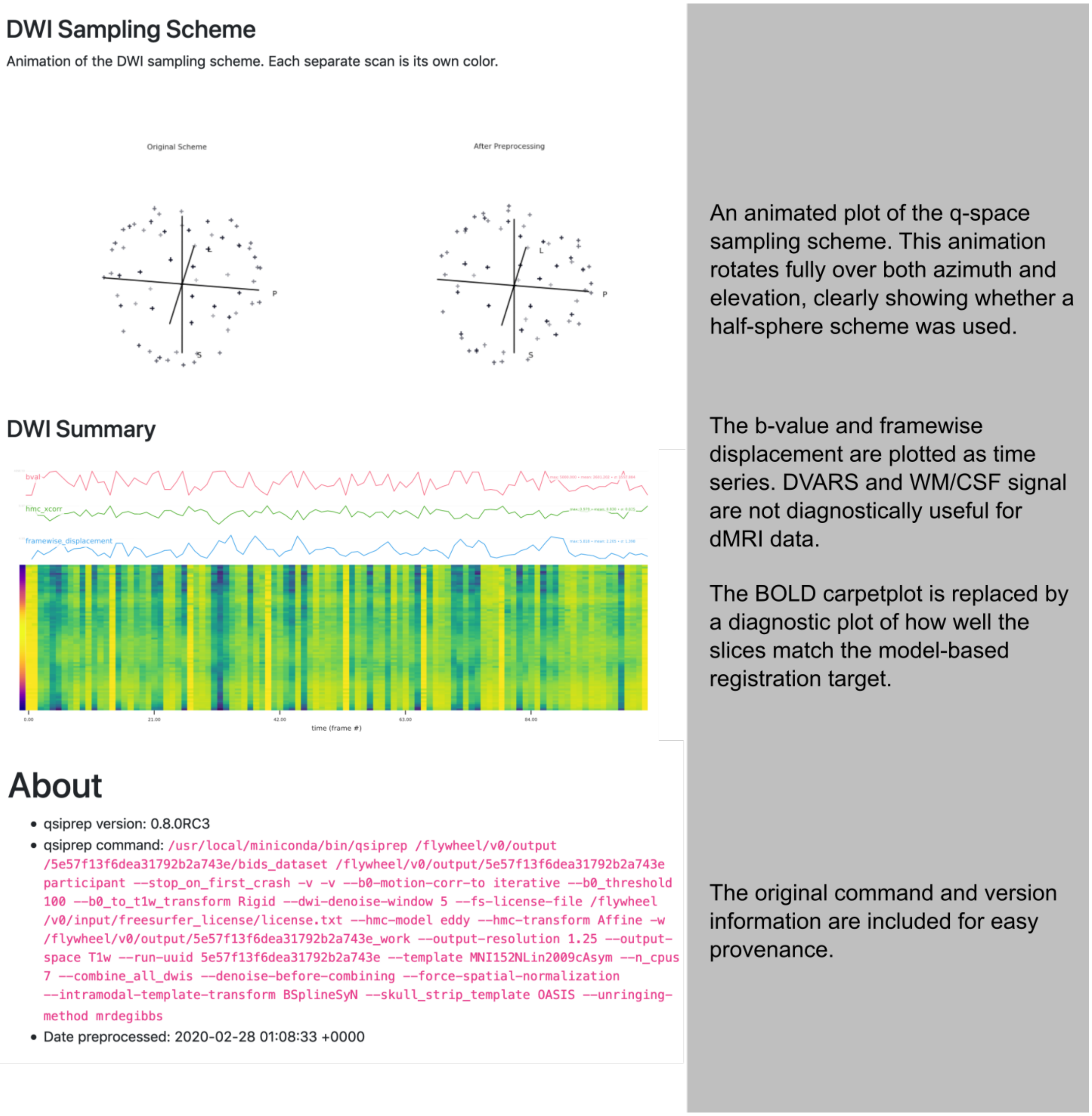


**Fig S1 | QSIPrep preprocessing report.** An example report is displayed (left) along with descriptions of what each section represents and how to interpret its contents (right, in gray).

| Actual report from QSIPrep | Description of report content |
|---|---|
| **Dipy**<br>**MAP(L)MRI**<br>Directionally color-coded ODF peaks overlaid on the b=0 reference image.<br>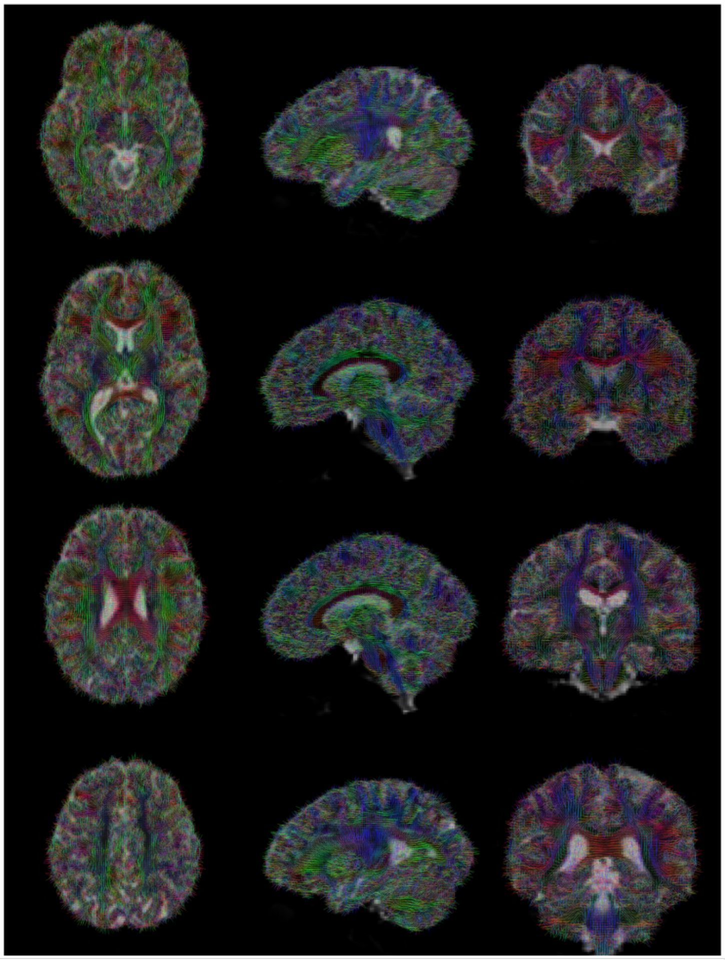 | A mosaic is shown of slices from the center of the brain. ODF or FOD peak directions are plotted as lines in each voxel.<br><br>This image is useful for quickly assessing if ODF peaks are pointing in the right direction. |

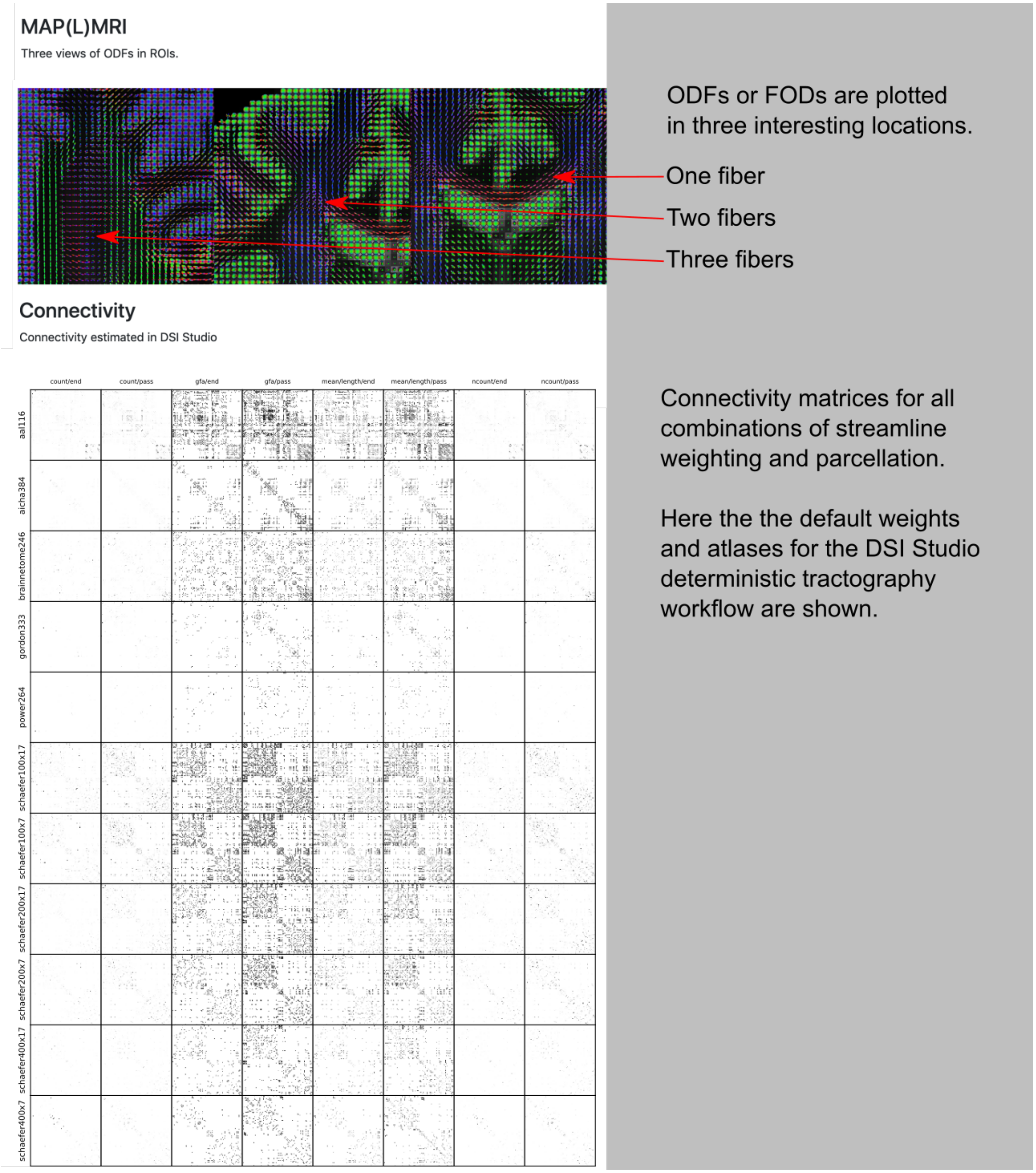


**Fig S2 | QSIPrep reconstruction report.** An example report is displayed (left) along with descriptions of what each section represents and how to interpret its contents (right, in gray).

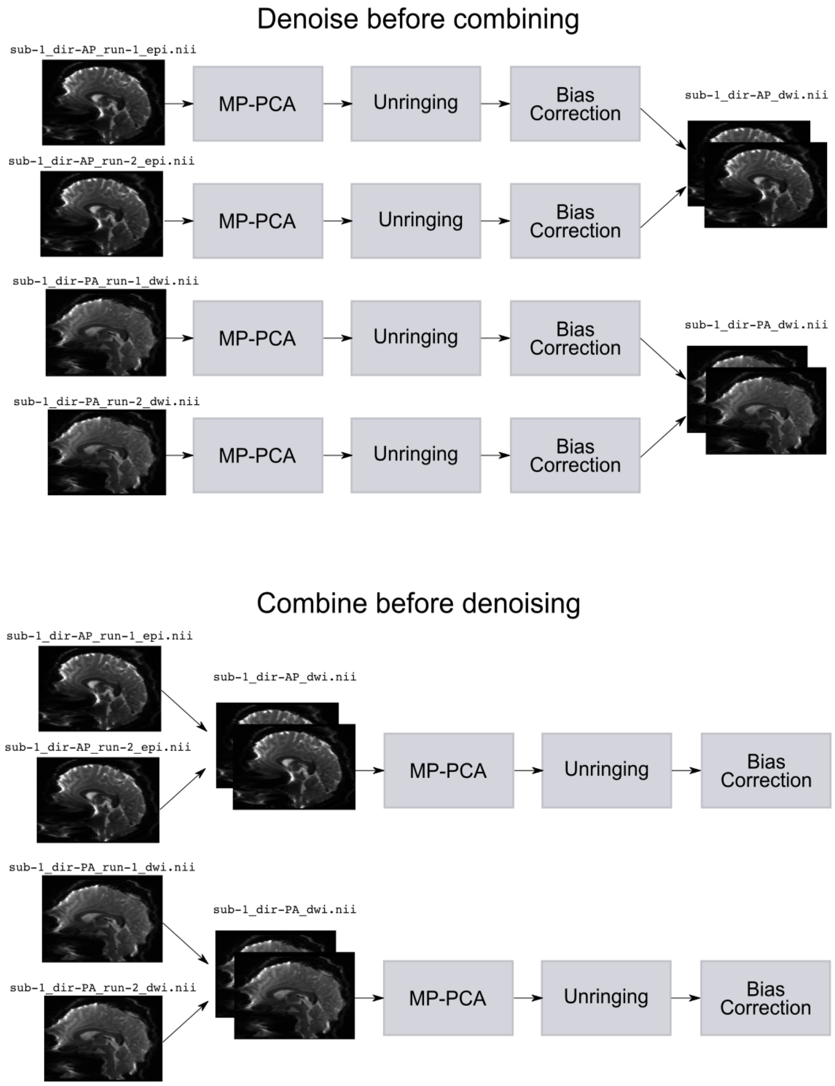


**Fig. S3 | Denoising workflows.** Two different workflows for denoising and combining multiple dMRI scans from a single session with multiple phase-encoding directions.

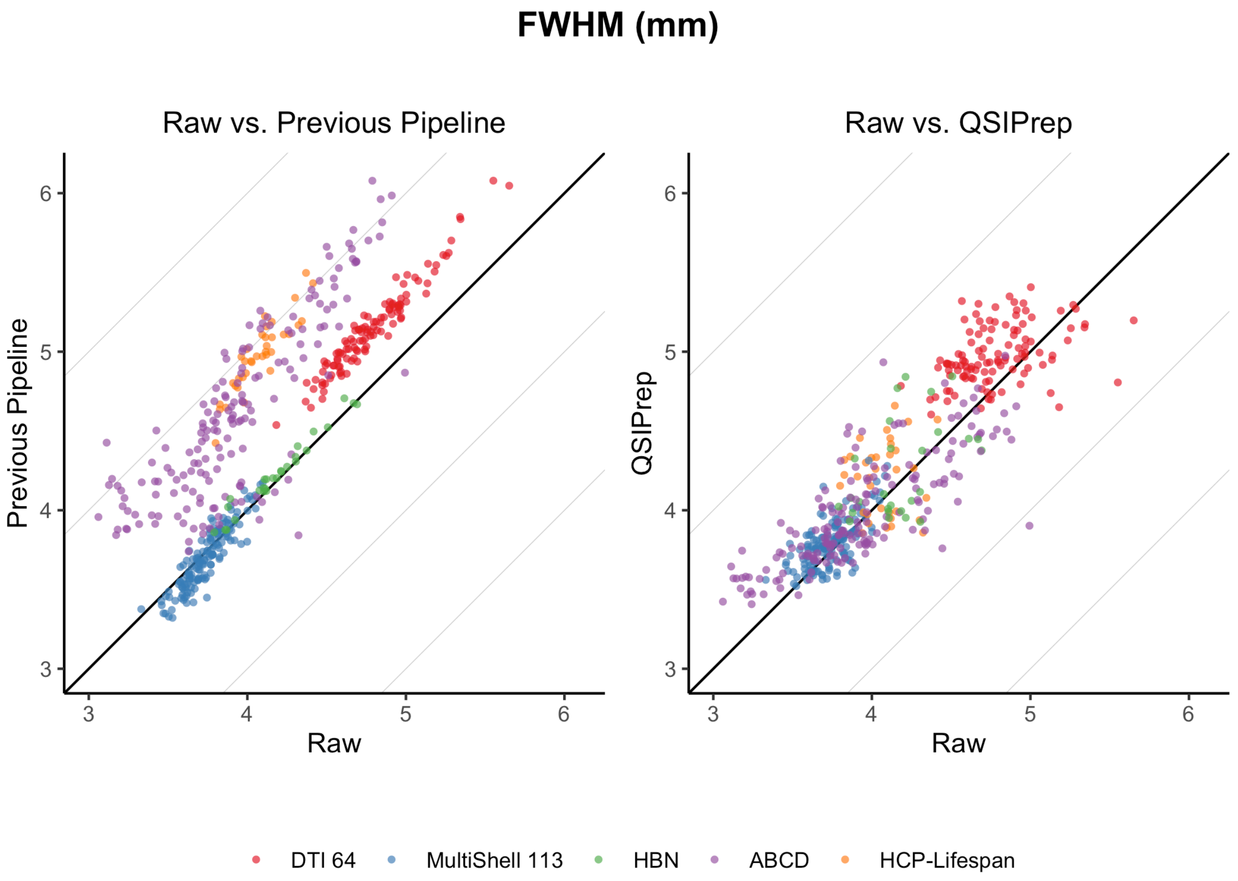


**Fig S4 | Comparing added smoothness from QSIPrep and previous pipelines.** Preprocessing generally increases the spatial smoothness of images relative to the raw images. Here the raw image smoothness (x-axis) is compared to the same images after being processed by the published pipeline for each dataset (left) and QSIPrep (right). Preprocessing using either pipeline increases smoothness relative to the raw images, but QSIPrep introduces far less additional smoothing. The direct comparison between QSIPrep and the Previous Pipeline is presented in **Fig 2**.

**Supplementary Note 1: dMRI Reconstruction overview**

*Sampling schemes and reconstruction methods*

Diffusion MRI signal is related to the water diffusion process in each voxel. Diffusion can be sampled along a specific direction by manipulating the magnetic field gradients, while the signal's sensitivity to water diffusion can be manipulated by altering the gradient strength. Although directions are typically described as unit *b*-vectors and gradient strengths are described in *b*-values in units of s/mm$^2$, each dMRI image is a sampling of coordinate $\boldsymbol{q}$ in *q*-space[1] where

$$\boldsymbol{q} = 2\pi\boldsymbol{u}\sqrt{3b/\tau}\,,$$

for unit *b*-vector $\boldsymbol{u}$, *b*-value $b$, and duration $\tau$ (which is typically the difference between pulse separation and duration parameters $\Delta$ and $\delta$) respectively. This relationship is the source of the name QSIPrep, as it can process any single diffusion encoding *q*-space image set.

It is useful to disambiguate a *sampling scheme* from a *reconstruction method,* as these often confusingly share the same names. The set of coordinates sampled in a dMRI scan constitute the *q*-space sampling scheme. When the samples are on the surface of a sphere (typically referred to as the "shell") is called a single-shell acquisition, or often colloquially referred to as a DTI because it is commonly used as input to the DTI reconstruction method. Multiple concentric shells are known as multi-shell. Sampling *q*-space on a set of Cartesian coordinates is commonly known as a Diffusion Spectrum Imaging (DSI) sequence[2]. A random set of coordinates in *q*-space can be used for a compressed-sensing DSI[3–5] (CS-DSI).

A *reconstruction method* is an equation or algorithm that uses the dMRI signal acquired on the sampling scheme to characterize the water diffusion process. For example, the Diffusion

Tensor Imaging reconstruction method is a least-squares fit of the dMRI signal to a 6-parameter tensor model[6]. Similarly, one can perform a Constrained Spherical Deconvolution (CSD) on shelled sampling schemes because their $q$-space coordinates can be defined on $S^2$, the same domain as the spherical harmonic basis set[7,8]. Other reconstruction methods model the full 3D $q$-space such as the simple harmonic oscillator reconstruction and estimation (3dSHORE)[9] and mean apparent propagator MRI (MAPMRI)[10]. These methods can be applied to any sampling scheme that has more than one unique shell.

The output of a reconstruction method is typically an *orientation distribution function (ODF)*, which maps directional unit vectors to a magnitude that reflects the diffusivity of water in that direction (a diffusion ODF) or the density of fibers in that direction (a fiber ODF). Methods like generalized $q$-sampling imaging (GQI)[11], 3dSHORE, and MAPMRI estimate a diffusion ODF while CSD estimates a fiber ODF.

While diffusion and fiber ODFs measure different phenomena, both should exhibit local maxima aligned with or close to the directions of fiber populations in each voxel. In this sense both are appropriate inputs for deterministic tractography, which is why all ODF outputs from QSIPrep reconstruction workflows produce DSI Studio “fib” files in addition to the native file format of the software used to reconstruct them.

*Software implementations*

The three main software packages used for ODF reconstruction in QSIPrep are DSI Studio[12], MRtrix3[13], and DIPY[14]. Each software package has its own file format and orientation conventions, and each provides complimentary features. DIPY has an extensive library of reconstruction methods and a robust visualization system used in QSIPrep’s reports **(Figure S3)**.

MRtrix has a toolchain highly optimized for single and multi-shell CSD. QSIPrep also includes the MRtrix3Tissue[15] fork of MRtrix3 so that both single and multi-shell sampling schemes can benefit from either SS3T-CSD or MSMT-CSD. DSI Studio provides a feature-rich visualization tool that enables real-time deterministic tractography, ODF and surface visualization, and a number of quality-control tools.

**Supplementary Note 1: References**


1. Callaghan, P. T. *Principles of nuclear magnetic resonance microscopy*. (Oxford University Press on Demand, 1993).

15. MRtrix3Tissue | MRtrix3Tissue is a fork of MRtrix3. https://3tissue.github.io/.

**Supplementary Note 2: Automatically generated methods boilerplate examples**

*2.1 Preprocessing example:*

Preprocessing was performed using *QSIPrep* 0.9.0beta1, which is based on *Nipype* 1.5.0 (Gorgolewski *et al*. (2011); Gorgolewski *et al*. (2018); RRID:SCR_002502).

**Anatomical data preprocessing**

The T1-weighted (T1w) image was corrected for intensity non-uniformity (INU) using N4BiasFieldCorrection (Tustison *et al*. 2010, ANTs 2.3.1), and used as T1w-reference throughout the workflow. The T1w-reference was then skull-stripped using antsBrainExtraction.sh (ANTs 2.3.1), and using OASIS as target template. Spatial normalization to the ICBM 152 Nonlinear Asymmetrical template version 2009c (Fonov *et al*. 2009, RRID:SCR_008796) was performed through nonlinear registration with antsRegistration (ANTs 2.3.1, RRID:SCR_004757, Avants *et al*. 2008), using brain-extracted versions of both T1w volume and template. Brain tissue segmentation of cerebrospinal fluid (CSF), white-matter (WM), and gray-matter (GM) was performed on the brain-extracted T1w using FAST (FSL 6.0.3:b862cdd5, RRID:SCR_002823, Zhang, Brady, and Smith 2001).

**Diffusion data preprocessing**

Images were grouped into two phase encoding polarity groups. Both groups were then merged into a single file, as required for the FSL workflows. Several confounding time-series were calculated based on the preprocessed DWI: framewise displacement (FD) using the implementation in Nipype (following the definitions by Power *et al*. 2014). The head-motion estimates calculated in the correction step were also placed within the corresponding confounds file. Slicewise cross correlation was also calculated. The DWI time-series were resampled to ACPC, generating a preprocessed DWI run in ACPC space.

Many internal operations of qsiprep use Nilearn 0.6.2 (Abraham *et al.* 2014, RRID:SCR_001362) and DIPY (Garyfallidis et al. 2014). For more details of the pipeline, see the section corresponding to workflows in qsiprep's documentation.

Deconvolution: Validation Using Diffusion-Weighted Imaging Phantom Data." *Neuroimage* 42 (2). Elsevier: 617–25.
